# Single-cell neural network classifiers reveal that PM21 NK cell expansion is dependent on B cell signaling

**DOI:** 10.1101/2025.09.23.676816

**Authors:** Michael Tsabar, Shaoline Sheppard, Sarah Sturtevant, Yinyin Huang, Ryan Genga, Margaret Magaletta, Matthew Kodrasov, Elizabeth Tran, Kyle Smith, Juhyung Jung, Matthew Sullivan, Andre Kurlovs, Virginia Savova, Diana Stoycheva, Bruno Figueroa, Dennis Goehlsdorf, Alexandra Grella, Emanuele de Rinaldis, Giorgio Gaglia

## Abstract

In the field drug development ML/AI methods are being applied to improve drug production speed, costs, and reliability. In allogenic NK cell therapy production, one of the biggest challenges is the inherent variability in the donors that provide the starting material for NK cell expansion. In this study we performed PM21-mediated NK cell expansion on 26 donors, and in parallel performed single-cell transcriptomics on the same donor sample prior to expansion. Canonical differential expression analysis and cell state abundance did not highlight any significant difference between donors with high and low NK cell expansion yield. Instead, training neural networks classifiers for high-yield donors enabled identifying several highly predictive models with perfect cross-validation recall. Further investigation of the most predictive models unveiled a previously unknown role for B cell in supportive NK cell expansion. Overall, this study represents a blueprint for combining deep phenotyping and machine learning methods to unveil novel biology and improve the quality and speed of delivery of cell therapeutics to patients.

## INTRODUCTION

Immunotherapy has revolutionized the treatment of cancer providing improved efficacy and reduced toxicity compared to chemotherapeutic approaches. Natural Killer (NK) cell-based therapies are one promising form of immunotherapy, specifically because lower immunogenicity of NK cells renders them suitable to allogenic application. For this reason, NK cell-based drug products can be derived from healthy donor peripheral blood mononuclear cell (PBMC) collections, allowing for an off-the-shelf treatment approach while lowering the incidences of adverse side effects (e.g. graft-versus-host disease) in contrast to T cell therapies [1–5]. Even though NK cells do not inherently possess the expansion potential necessary to produce commercial-scale quantities, recently engineered PM21 nanoparticles (“PM21”) trigger the ex vivo expansion of PBMC-derived NK cells, circumventing this problem [6, 7]. PM21 are derived from the plasma membranes of K562 cells overexpressing membrane-bound IL-21 and 4-1BBL. Compared to conventional feeder-cell expansion, PM21 eliminate the risk of feeder cell carryover contamination in the drug product and induce a hyperfunctional cell state [8].

The process of NK cell expansion for allogenic therapy is a multi-step process [9–14]. Blood is collected from a CMV-positive donor, subject to leukapheresis to isolate PBMC’s, which are then depleted of CD3+ cells by negative selection to arrive at a “starter culture”. The latter is then subject to two PM21 stimulation, which cause the expansion of NK cells resulting in a clinical product currently being tested in trials for two types of blood cancer: recurrent, refractory acute myeloid leukemia (R/R AML, NCT04220684 and NCT05712278) and high-risk myeloid malignancies undergoing allogeneic hematopoietic stem cell transplant (HSCT; NCT05115630 and NCT05726682). However, the success rate of manufacturing campaigns expanding NK cell ex vivo with PM21 starting from PBMC’s is only around 50% and even in successful campaigns, NK cells PM21 expansion rates are highly variable donor-to-donor [1, 11]. The source of NK cell expansion variability does not solely depend on the donor’s NK cell phenotype; PBMC’s are a frequent source of NK cells for clinical applications [6, 15–18] and cellular communication from other cell types has been demonstrated to effect NK cell expansion and differentiation [19–23]. The causes of the expansion heterogeneity are still not understood and a biomarker for donor expansion potential is not available. Lack of robustness and predictability increases production cost severely limiting the manufacturability and applicability of allogeneic NK cell therapy [18, 24, 25].

Since other cell types can influence both activation and suppression of NK cell growth and function, understanding the full profile of the starting material might provide insight in the variability of PM21 expansion as a reliable predictor of their NK cell capacity for expansion. In the last decade single cell RNA transcriptomics has become an extremely powerful tool to achieve a detailed understanding of heterogenous populations of cells, enabling identification of novel cell-types, gene expression dysregulation across conditions and prediction of developmental trajectories. Single-cell RNA sequencing (scRNA-seq) datasets are extremely rich, providing quantitative expression for thousands of genes from millions of individual cells. However, the transcriptome of different individuals can be highly variable, confounding the detection of true condition-relevant patterns. Moreover, the overwhelming size of data often results in only a fraction of this resource being utilized and translated into clinically relevant knowledge and applications. To harness the full potential of such data, machine learning approaches have become an attractive tool in bioinformatics research, integrating biological knowledge with computational techniques to extract relevant biological insights. The use of machine learning in single cell research provides unprecedented opportunities to enhance personalized medicine, reduce treatment costs and positively impact overall population health.

In this study, we investigate the sources of PM21 NK cell expansion variability by deeply profiling the starter cultures before PM21. We generated scRNA-seq of CD3-depleted PBMC’s from 26 donors before a 12-day PM21 stimulation protocol, during which we measured NK cell counts. Since we gained limited insights through conventional scRNA-seq analyses, we employed a machine learning approach leveraging the convolutional neural network ScaiVision based on CellCNN [26] to determine predictors of strong expansion potential in response to PM21 stimulation. We were able to successfully develop models predicting donors with high-yield NK cell expansion potential (AUC = 0.89), using only pre-PM21 information. Surprisingly, the cell type with the highest predictive value were activated B cells, rather than NK or innate immunity cells. Additionally, using integrated gradient method, we were able to extract the genes driving the prediction and distill it into two 100-gene signatures, maintaining the original predictive performance. The ability to derive novel insights and predictive gene sets from a small cohort of omics-profiled samples represents a translation framework for boosting drug discovery and clinical trial design.

## RESULTS

### PM21-stimulated NK cell expansion is supported by the presence of other cell populations and expansion rates vary by donor

To study the NK cells expansion process, we obtained and expanded individual donors using a PM21-based, 12-day NK cell expansion process (schematic in Fig. 1A). First, we tested whether the expansion variability was due to the presence of other non-NK cell types in the starter culture. We compared the same n = 4 donors expanded either from a T-cell depleted culture (as per clinical protocol) or from a purified CD56+ NK cell population. Seven days after PM21, the T-cell depleted cultures expanded 4-fold better than the pure NK cell population (Fig 1B). In both starting condition the expansion was highly variable (coefficient of variation = 0.57 and 0.87 respectively). Specifically, we observed that when culturing NK cells with no other cell type, a 10-times higher seeding concentration was required to achieve the same rates of expansion as T-cell depleted PBMCs (n=1, Fig S1A).

**Figure 1:**
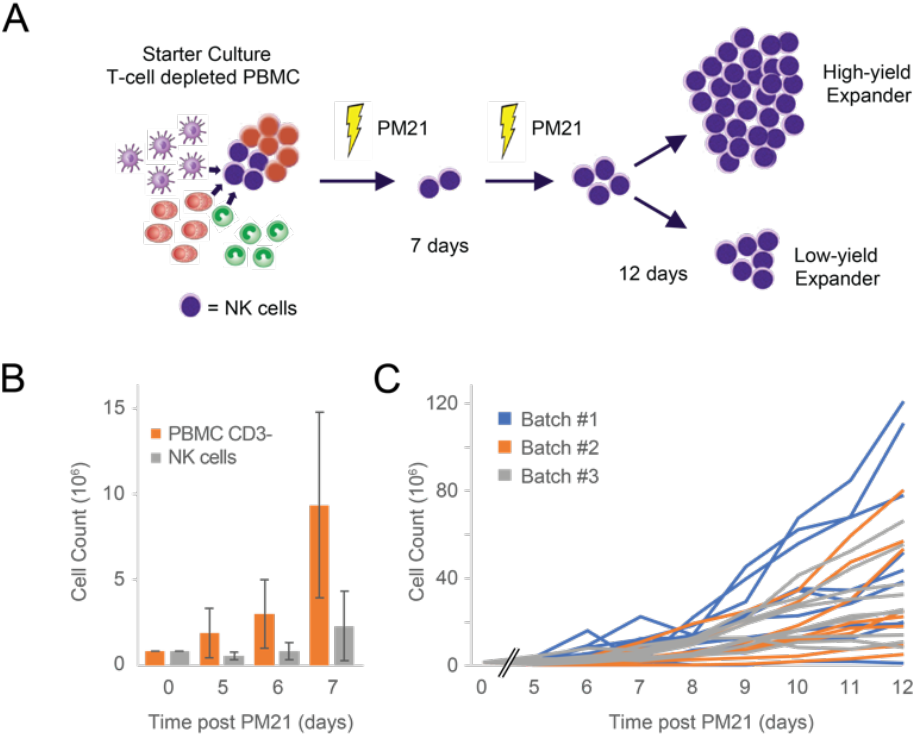
**A**. Expansion protocol graphical summary. **B**. Comparison of expansion of T-cell depleted starter cultures (CD3-PBMC, orange) and purified CD56+ NK cells (NK cells, grey bar) from the matched donors. Mean and standard deviation, n=4. **C**. NK cells expansion dynamics for starter culture from n=26 donors. Measurements are taken every day starting from day 5 after the first PM21 stimulation.

We next expanded T-cell depleted PBMC samples from n = 26 donors in three batches. After a 12-day expansion protocol the final NK cell counts ranged from 1 to 120 million NK cells, with the average being 40 million (Fig. 1C). Despite the increased number of samples, the expansion variability remained high (CV = 0.79) and this was observed to be independent on the batch of the expansion (Fig S1B,C). Finally, we tracked over time the correlation of NK cell counts with final concentration at the end of process, to determine when the final concentration of NK cells could be accurately predicted; we observed that correlation increased gradually over time and reached 90% only at Day 10 (Fig S1D). These results demonstrate that the process of NK cell expansion is highly variable and unpredictable, and that non-autonomous interactions between NK cells and other cell populations in the culture are required for successful expansion.

### Cell proportion and differential gene expression analysis on cells prior to expansion fail to define clear key features of high-yield donor

In order to understand the molecular basis of the variability in donor-to-donor expansion we performed single cell transcriptomics on the 26-sample cohort which showed high level of PM21-expansion variability (Fig 1B). Frozen samples prior to expansion (day 0) were thawed, single-cell suspension prepped and sequenced both for transcriptomics and surface antigen detection (CITE-seq). The sequencing was analyzed through standard bioinformatics pipeline (CellBridge [27]) with a total of n=210,183 single cells annotated using the SignacX algorithm [28] (Fig 2A). The major cell types expected were retrieved in proportions congruent with prior knowledge and marker review confirmed the quality of the cell type annotation (Fig S2A). As expected, given the anti-CD3 negative selection performed prior to cryopreservation, the T-cell compartment was minimally observed in these samples, with the dominant populations being myeloid (monocytes and dendritic cells) and B cells (Fig 2B).

**Figure 2:**
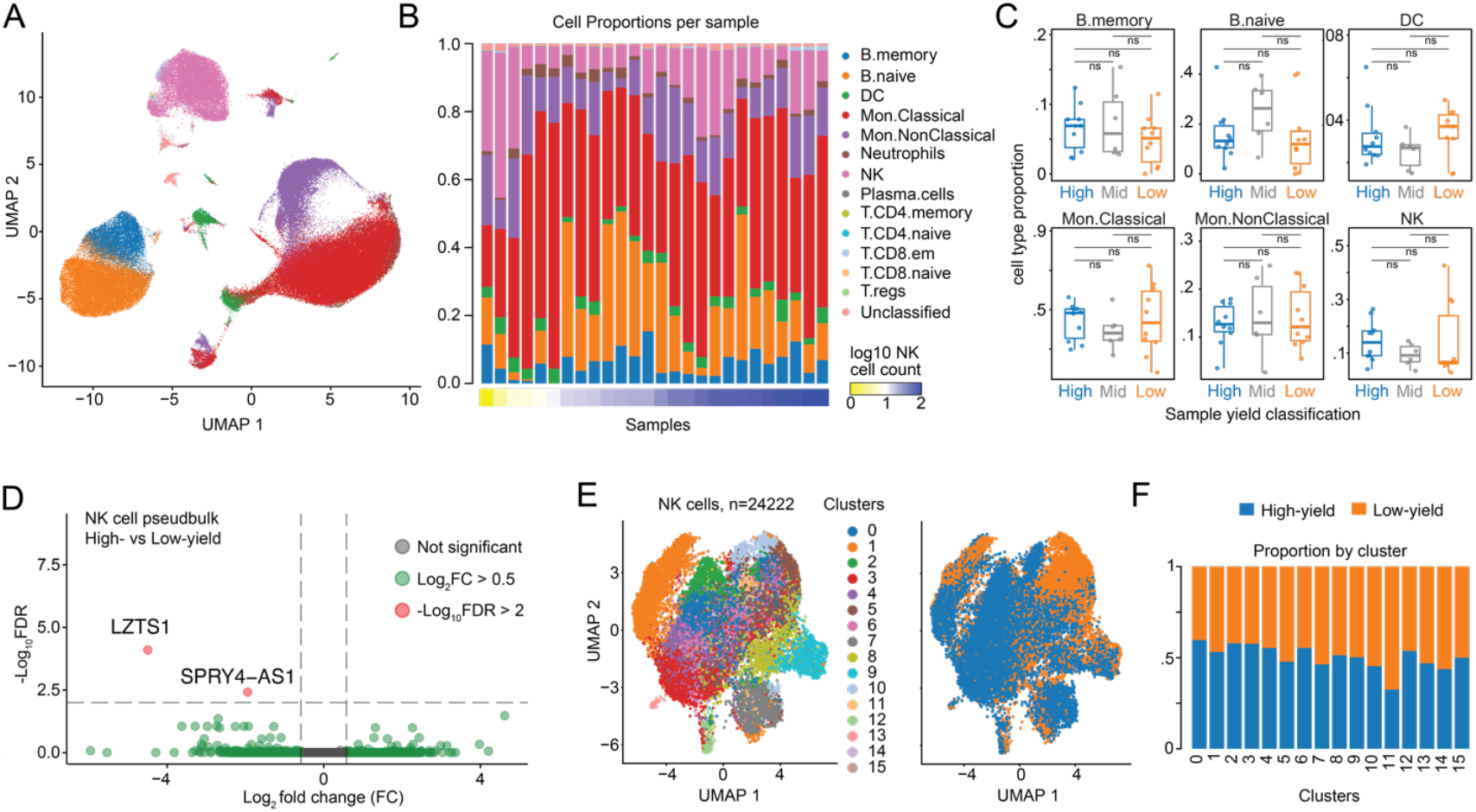
**A**. UMAP of scRNA-seq data (SignacX cell states, n = 210,183 cells). **B**. Distribution of cell states by samples ordered by expansion rate (n=26 samples, normalized by total cells per sample). **C**. Box plot with whiskers of SignacX cell state proportions subdivided by high-, medium- and low-yield expanders (High, n=10, blue, Mid n=6, grey, Low n=10, orange). **D**. Volcano plot of pseudobulk differential gene expression of NK cells in high-versus low-yield expanders. **E**. UMAP of NK cells from high- and low-yield expanders (n=24,222 cells) colored by Louvain cluster or by yield group. **F**. Proportion of expansion group per Louvain cluster.

We then separated the samples in three groups based on the endpoint expansion success rate - 10 high-yield expanders, 10 low-yield expanders and 6 medium-yield expanders – and asked whether the cell-level features are causing the difference in expansion rates. The cell-type proportions per sample were not found to be different between the expansion groups for either NK cells or any other major cell type (Fig 2C and S2B). Next, we tested whether there were differences in the phenotypes of cells, rather than the number of cells. Comparing high-versus low-yield expanders, the differential expression pseudobulk analysis highlighted only two genes significantly dysregulated in NK cells, none of which has any relevance on NK cell biology (Fig 2D). Similar cell-state specific differential expression analysis found limited phenotypic differences in all cell types surveyed (Fig S2C). Finally, we explored whether the expansion differences could be due to a phenotypically distinct subset of NK cells highly responsive to PM21. We in silico isolated NK cells from all donors, re-clustered them by transcriptomics similarity, and calculated whether any of the clusters was more abundant in high- or low-yield expanders (Fig 2E, F). None of the clusters was skewed towards either population suggesting that the sub-states of NK cell was evenly distributed. Considering these lines of evidence together, canonical analyses failed to highlight any actionable molecular insight that could explain the expansion variability.

### Shallow neural net modeling predicts expansion success from-single cell transcriptomics data

Since the canonical comparison of cell numbers and phenotypes failed to explain the expansion heterogeneity between samples, we employed ScaiVision (Scailyte AG), a supervised representation learning approach using shallow neural networks (Fig 3A) to predict sample-level NK cell expansion outcomes. As input, the model takes single-cell data (transcriptomics or CITE-seq), using the sample’s expansion yield as the training label. The input data (a |genes| x |cells| matrix) is first dimensionality-reduced via PCA (Step 1, PCA), using a pre-computed transformation matrix (W^d^) which is not altered during training. Thereby, the number of trainable parameters is reduced. Subsequently, the data is subsampled to generate multi-cell inputs of consistent size (Step 2, Sampling), effectively augmenting the data and mitigating the limitations of a small sample size. The augmented data is then passed through a linear layer (defined by weight matrix W^f^ and bias values b^f^). The columns of this matrix can be interpreted as ‘filters’ which are applied to each input cell, thereby generating cell response values (Step 3, Filter application). The values are then clipped to non-negative values and pooled across the top k responding cells for each filter (Step 4, Pooling). This special pooling process allows to focus on the most relevant cell populations while reducing the influence of outliers. Finally, a second linear layer (defined by weight matrix W° and bias b°), combined with a SoftMax function, produces sample-level probabilities for high or low expansion (Step 5, Output Calculation).

**Figure 3:**
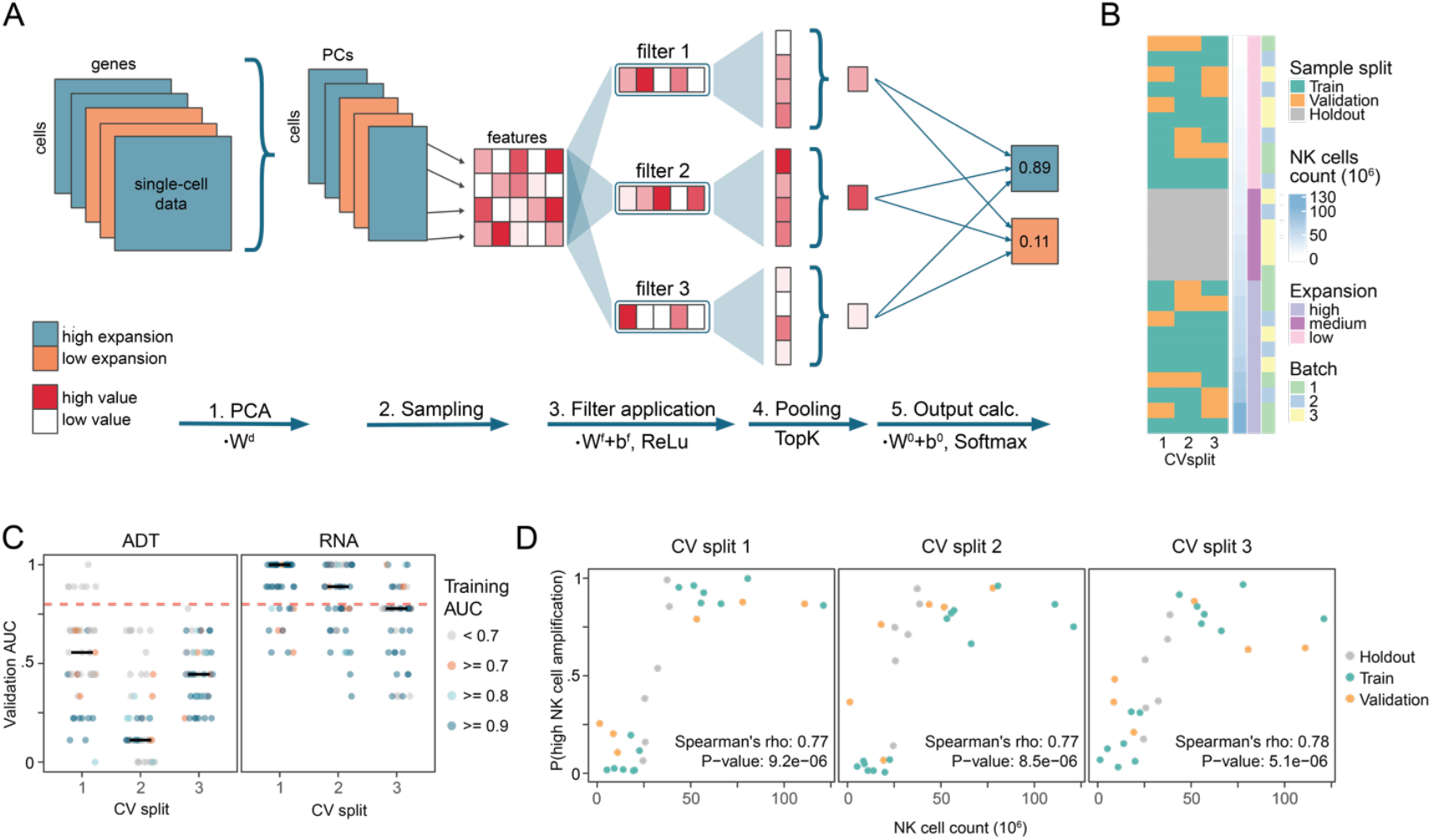
**A**. Graphical representation of the ScaiVision algorithmic flow and network topology. The input data, provided as a matrix of size |genes| x |cells| is reduced in dimensionality by a linear transformation using the dimension reduction matrix W^d^. This matrix is pre-computed and not optimized during training, which reduces the number of parameters that need to be optimized during training. In a second step, the reduced data matrix is passed through a linear layer, defined by the weight matrix W^f^ and the bias b^f^. Not shown is the reduction of values to positive values. The weight matrix W^f^ is symbolically shown in the right corner: for each filter it contains one column vector of weights, defining the specificity of each filter in feature space. Filtered cell responses are pooled by averaging across the top k responses. Predictions are finally computed by a second linear layer (defined by weight matrix W° and bias b°), followed by a softmax operation, which yields values in the range [0,1]. The final outputs can be interpreted as probabilities for a high or a low expansion. **B**. Cross-validation split (CV-split) division of n=26 samples into training (green, n=14), validation (n=6, orange) and holdout sets (n=6, grey, mid-yield samples). **C**. Area under the curve (AUC) from validation of n=50 models per CV-split for models restricted to ADT (left) or RNA data(right). Color represents AUC achieved by each model in the training phase. Horizontal black line represents median AUC. **D**. Scatter plots of the probability of classification as “high yield” by top 5 performing models against the observed number of NK cells at day 12 of PM21 protocol. Color represents whether the sample was used to train, validate, or as part of the hold-out group. Indicated spearman correlations rho and p-values were calculated using all samples.

To enhance the likelihood of identifying high-performing and generalizable models for downstream analysis, we employed a Monte Carlo cross-validation strategy to evaluate model performance. From the high- and low-yield expander samples, we created 3 cross-validation splits (CV-splits), each splitting the 20 samples into 14 training and 6 validation samples (Fig 3B, Table S1). For each CV-split, 50 ScaiVision models were trained, each using a different set of hyperparameters. Each of the 50 runs were evaluated with the area under the curve (AUC) metric for the ability to predict training and validation sample sets.

Models were first trained using the entire CITE-seq dataset encompassing both RNA and ADT modalities (FigS3A). High performing ScaiVision models were generated with 2 out of 3 folds reaching a median AUC above 0.77. To investigate if the performance was driven by one of the modalities, we further trained models using RNA and ADT data separately. When the ScaiVision models were trained using the ADT data a handful of models reached AUC > 0.8 on the training data (Fig 3C left, blue and cyan dots), but these models consistently fail to perform better than random in the validation set (AUC < 0.5). Of note, since each of these training runs is based on an individual parameter initialization and hyperparameter setup, it is expected that many training runs fail, mostly due to non-optimal random initialization of the filters. However, no model trained on the ADT data achieved AUC > 0.8 for two of the cross-validation (CV) splits, which means that under no circumstances the classifier generalized to unseen samples. Instead, using only the single cell RNA data, the training yielded multiple high-performing models for each of the CV-splits (Fig 3C right), outperforming the performance achieved when using the RNA and ADT combined. Models with AUC > 0.8 on both training and validation sets accounted for 39% of all models (70%, 32%, 14% for CV-split 1, 2, and 3 respectively). In addition, multiple models in every CV-split were able to predict unseen samples with perfect accuracy (AUC = 1). These performance results are indicative of a clear discriminative signal present in the single-cell transcriptomics data.

To determine whether the output probabilities generated by the models accurately reflected the underlying variability of the sample’s expansion potential, we analyzed the correlation between the two. We filtered models for those with a training and validation accuracy of at least 80% and selected the 5 models with lowest cross-entropy loss on the validation data for each fold (except for CV split 2, where only 4 models passed the criteria). For all CV-splits we observed a clear relationship between the NK cell count after expansion and the mean predicted probability of belonging to the high-yield class (Fig 3D, green and orange dots), as indicated by the Spearman correlation on all samples (Spearman correlations = 0.77, 0.77, 0.78, and p-values = 9.2e-06, 8.5e-06, 5.1e-06). Additionally, we analyzed the held-out set of samples with intermediate NK cell expansion yield, separately from the training and validation samples (mid-yield n=6). While it would be expected that the trained models would struggle to sort these intermediate and potentially noisy measurements, the mean probability outputted by the models was highly and significantly correlated with the final NK cell numbers (Spearman correlations: 0.89, 0.77, 0.83, p-values 0.03, 0.10, 0.06). In summary, the ScaiVison method trained on pre-PM21 transcriptomics data was able to create multiple models with perfect ability to classify unseen samples and the probabilities for high expansion predicted by the top models quantitatively tracked the number of NK cells obtained by the protocol.

### Interpretation of top performing models suggests that B cells are the cell type most predictive of expansion success

To acquire a deeper understanding of the underlying biological mechanisms driving the prediction of our models, cells selected by the highest performing models were examined. This process is facilitated by the shallow architecture of the neural networks underlying ScaiVision. Cells in the top 3% based on ScaiVision cell filter scores were selected and plotted (Fig 4A). Surprisingly, the cell-type distributions in both CV-splits 1 and 2 show clear enrichment of B cells, which had not been previously linked to PM21-driven NK cell expansion in the literature (Fig 4B, Table S2). Differently from the other CV splits, CV-split 3 distribution of highly predictive cells is much more similar to that of the overall cell population, with a present but only slight enrichment of B cells. The difference in cell preference can in fact account for the generally lower performance of trained models in CV-split 3 (Fig 3C). Among the 6 validation samples used for CV-split 3, two samples have an extremely low percentage of B cells (3.29 %, 1.4 %), not representative of the average cell population (median B cell fraction is 22%). This may have impacted the model’s ability to learn to identify the most predictive cells and accurately predict the NK cell expansion in CV-split 3.

**Figure 4:**
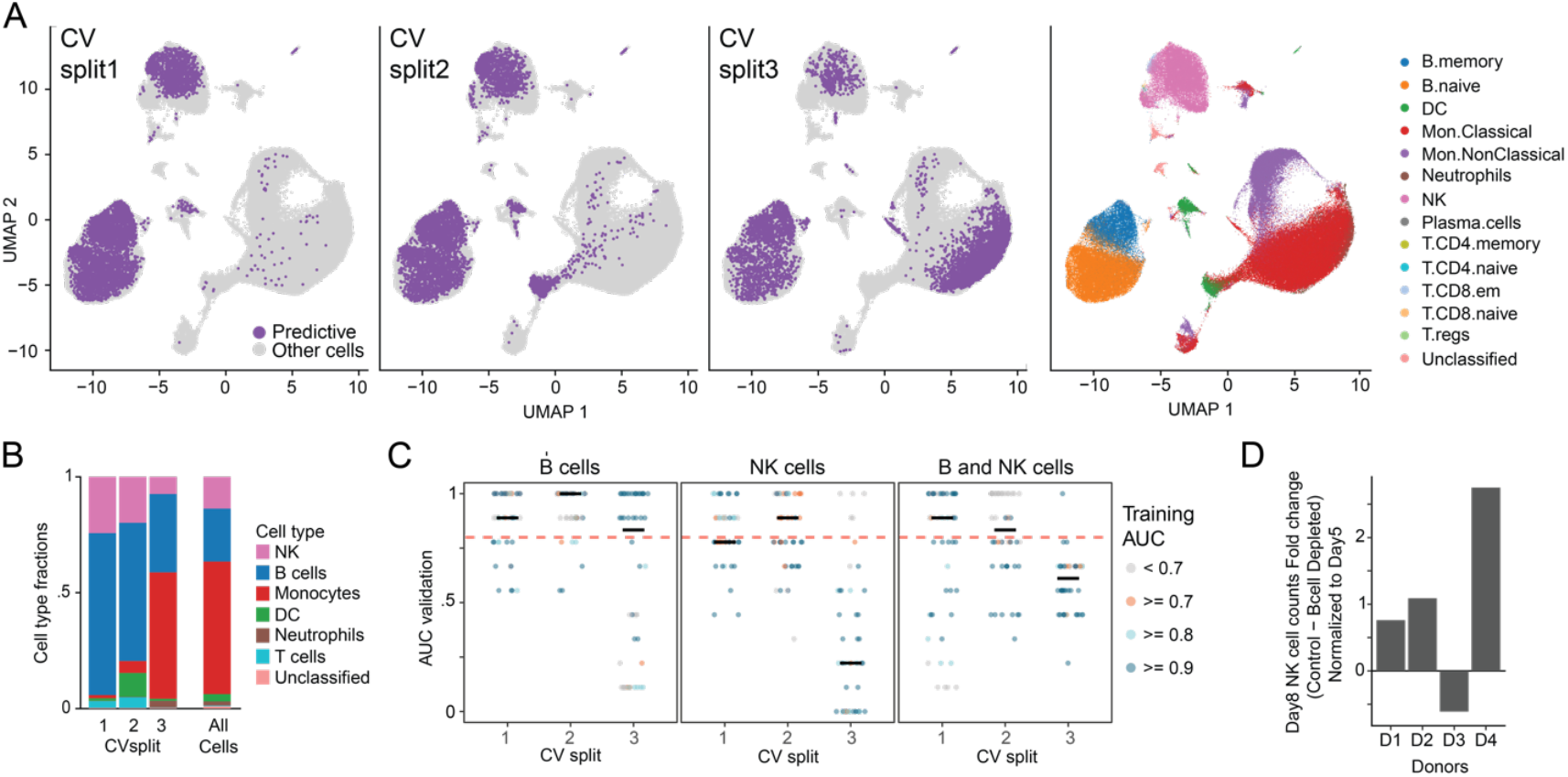
**A**. UMAP of scRNA-seq data labelled by predictive cell call by ScaiVision (left, purple) or by SignacX cell states (right). **B**. Proportion of predictive cells (left) or all cells (right) labelled by SignacX cell states. **C**. Area under the curve (AUC) from validation of n=50 models per CV-split. Models were trained only using B cells (left), NK cells (middle) or B and NK cells (right). Color represents AUC achieved by each model in the training phase. The horizontal black line represents median AUC. **D**. Fold change difference of NK cell counts at day 8 between control and B-cell depleted (normalized to day 5). n=4 donors.

The finding that B cells were the most consistent cell-type selected by our models prompted us to investigate if a better performance could be achieved when focusing on these cells alone. We therefore trained ScaiVision models using either only B cells, only NK cells, or B and NK cells together. The B-cell only training led to a staggering majority of models with AUC > 0.8 (Fig 4C). These models outperformed NK cells and combined B and NK cell models. The majority of models in both only B-cells and all-cells setups reached AUC > 0.8 (70 % vs 51%, Fig. 4C left and 3C). Models trained with either NK cells or B and NK cells performed worse than B cells alone (Fig 4C center and right). This indicates that most of the signal driving model training in the all-cells-setup was retained when selecting for B-cells only and that additional use of NK cells did not add information.

To experimentally test the relevance of B-cells in the expansion process, we compared the ability of 4 donors to expand in presence or absence of B cells. We ran PM21 expansion reactions in four donors, where in addition to the negative selection with anti-CD3 we added anti-CD19 selection. Hence each donor sample was either depleted of T cells, as in the standard PM21 protocol, or depleted of both T and B cells. In 3 out of the 4 four donors we observed a worse performance of B-cell depleted samples (Fig 4D).

### Extraction of predictive genes enables reducing classifier to interpretable minimal models

We next leveraged the models derived by ScaiVision to investigate novel biology of NK cell expansion, by first identifying the genes responsible for driving performance in the high accuracy models and then generating reduced models. For each of the B-cell and all-cell setups, the top 9 models with high accuracy and lowest cross-entropy loss were selected (three models per CV split). The models were then analyzed using integrated gradients, to generate a gene-wise score for each cell selection configuration. The gene scores correspond to the contribution of each gene to the prediction of NK cell expansion potential. To determine whether the gene scores were representative of predictive power, we ran ScaiVision using the top 100 genes with the highest contribution score for both the B-cells and all cells experiments. High number of models were obtained, with an average validation AUC across CV-splits of around 0.87 and 0.88 for B and all cells respectively (Fig 5A & FigS3B). Moreover, the performance across CV-splits remained reasonably stable, where at least 50% of the models had a validation AUC > 0.75 for both signatures and all CV-splits. These observations still hold true for 2 out of 3 CV-splits when genes were subsetted to the B cells signature and training was performed using all cells (Fig 5A). Finally, we tested whether the top 100 gene sets from B-cells and all-cells model provided additive information by training models with the intersection of the two gene sets (35 genes) or the complementary sets (65 genes). In all scenarios, the performance of the models decreases (Fig S3C) This indicates that although both signatures share common predictive patterns, each captures distinct features that contribute to accurately predicting NK cell expansion potential. It should be noted that by restricting the training to a reduced number of genes, such as in the case of the top 100 genes above, models were constricted to learn from those genes that helped to achieve strong performance on the validation data in the original experiment. It is therefore expected that gene-restricted models achieve better performance than the PCA-based original models. The performance achieved in the original models is therefore a better estimator for how well models would generalize to further data. However, the high performance obtained for these gene-restricted experiments indicates that the selected genes, and their combined expression are the primary factors driving the performance in the original experiments.

**Figure 5:**
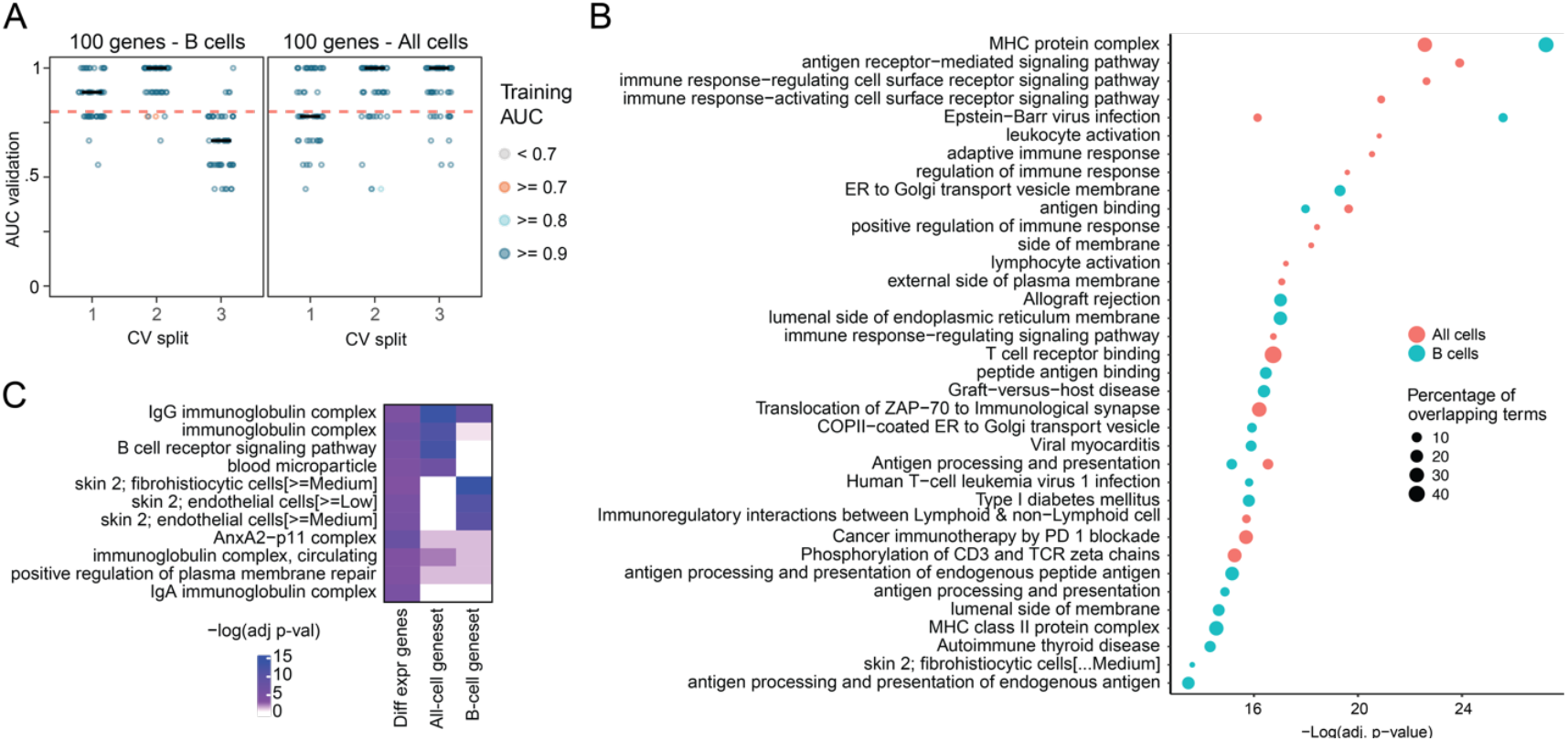
**A**. Area under the curve (AUC) from validation of n=50 models per CV-split. Models were trained only using the top 100 genes derived from B-cell models (left) or all-cell models (right) Color represents AUC achieved by each model in training phase. The horizontal black line represents median AUC. **B**. Gene set enrichment analysis (GSEA) of gene sets from B-cell models (green) or all-cells models (red). **C**. Comparison of GSEA p-values for all-cell and B-cell gene sets with gene sets derived from differential expression comparison between most predictive and least predictive B cells (top quartile of predictive ranking versus bottom quartile.

### Interpretation of ML models reproduces known biology and suggests new avenues of investigation

Both the cell-based models (B-cell and all-cell) and the reduced 100-gene models can provide insights into the biological pathways driving NK cell expansion. Hence, we examined the 100 gene sets as well as performing differential expression gene (DEG) analysis between predictive and non-predictive cells (highest quartile vs lowest quartile of predictive score distribution). Gene-set enrichment analysis suggest antigen processing, presentation and signaling is enriched in both top-100 gene sets (Fig 5B). Additionally, immunoglobulin complex genes are enriched in the top-100 gene sets and in the DEG comparison between predictive and non-predictive cells (Fig 5C).

The two 100-gene sets had 35 gene overlap which included a high number of genes related to antibody production, such as IGLC1, IGLC2, IGLC3. Additionally, immunoglobulins genes G (IGHG4 and IGHG3) were also found to be upregulated in predictive B cells as well as TNFRSF13B (TACI), whose signaling induces differentiation of B cells to plasma cells [29]. Despite the low numbers, plasma cells were found to be enriched in predictive cells (p-value adjusted through hypergeometric test < 0.05 across all 3 CV-splits, Table S2). Interestingly FCGR3A, a key receptor for antibody-dependent cellular cytotoxicity (ADCC) was among the genes present in the all-cell gene set. Combined with the increased antibody production of B cells, these results suggest that B cells could activate NK cells upon binding of this receptor to IGG coated targeted cells.

Among the 35 genes shared between the two 100-gene sets, eight genes code for MHC protein complex I (HLA-A, HLA-B, HLA-C) and II (HLA-DPB1, HLA-DQA1, HLA-DQB1, HLA-DRB1, HLA-DRB3). While increased expression of MHC class 2 on antigen presenting cells concords with B cell activation, the influence of expression changes in MHC I on NK cell expansion is less understood. Binding of NK cells receptors to MHC class I leads to inhibitory signals allowing self-tolerance [30]. This suggests that an upregulation of MHC I-coding genes could lead to a better survival and maintenance of both the NK cells and cells promoting the expansion of NK cells. Finally, genes related to cytokines (TNFAIP3), chemokines (CCL3, CCL5) as well as cytotoxic activity (GZMA, GZMH, NKG7, FGFBP2) [31] were present in all cell signatures. Heightened expression of these genes in NK cells suggests an activated cell state, which may enhance the survival and growth of NK cells. Altogether, these results agree with the current understanding of interactions between B and NK cells, suggesting potential explanations for the novel role for B cells in NK cell expansion. Importantly, these insights unveiled by machine learning trained models were not observed in canonical differential expression analyses and hence would not have been discovered using conventional single-cell data analytical approaches.

## DISCUSSION

Cell therapy is a promising therapeutic avenue for both oncological indications [5, 32] and autoimmune diseases [33–36]. Compared to T-cell therapy, allogenic NK cell-based therapy reduces the incidence of side-effects but remains an expensive therapeutic approach due to the high costs of production. Understanding the donor variability in NK cell expansion would allow pre-selecting high-yield donors, reduce unsuccessful expansion runs, and significantly lower production cost. In this work we applied single cell transcriptomics-based machine learning classification models to investigate the biology behind the donor-to-donor heterogeneity in NK cell expansion yields and we unveiled a previously unknown role of B cells in supporting NK cell expansion by PM21. Classification models based only on B cells, which hence have not seen NK or other cells, were able to predict high-yield NK cell expanders, just as well as models exposed to all cell types in the system.

Deeper phenotypic investigation of the single cell transcriptomic data highlighted antigen processing, display and peptide binding as the strongest pathway group associated with predictive B cells. The latter upregulate immunoglobulins and IGG, indicating that they could be in the process of plasma cell differentiation and antibody production. A well-known axis of NK-B cell interaction is antibody-dependent cellular cytotoxicity, where NK cells trigger apoptosis of opsonized cells and secrete IFNy, which promotes the immune response and consequently could support NK cell expansion. Hence, the B cell phenotypes might be mechanistically connected to the process of NK cell expansion. Alternatively, these B-cell phenotypes could be mechanistically unrelated to NK cell expansion and only serve as biomarkers for donors with “high-yield expansion potential” NK cells, for which this potential is not detectable by transcriptomics. Even though we were able to experimentally support that B-cell depletion is detrimental to NK-cell expansion with an intra-donor comparison, further work is required to conclusively disentangle these two hypotheses.

The heterogeneity in PM21-driven NK cell expansion is an example of a common biological conundrum: systems of cells that are phenotypically comparable at rest but react dynamically differently when exposed to the same stimulus. Despite exhibiting a range of 100-fold differences in NK cell expansion and applying scRNA-seq to profile the 26 starter cultures, conventional cellular analyses couldn’t provide insight into the drivers of expansion variability. A major limitation of these canonical single-cell analyses is the necessity of predefining groups of cells to compare (most often defined by cell type lineages and states, as classical and non-classical monocytes). Machine learning methods, as ScaiVision’s shallow neural classifier model, bypass this limitation and allow identification of cells and genes most associated with the desired outcome in an unbiased manner. When applied to the NK cell expansion data, ScaiVision analysis highlighted the novel role of B cells and generated a 100-gene signature able to infer the yield potential with high degree of precision; hence it both provided novel biological insights as well as generated relatively concise panel of genes predictive of high expansion.

More broadly, the drug development field faces other similar dynamical problems, where the relationship between starting point variability with endpoint outcome is crucial. One such fundamental problem is understanding the drivers of response to therapies, since in most instances cohorts of clinically similar patients display highly variable rates of drug response. The application of deep phenotypic characterization combined with unbiased methods for extracting predictive signals is a potential solution, and this work is a proof of concept for this approach. Importantly, here we demonstrate that combining single cell transcriptomics with shallow neural networks enables extracting meaningful signals from a relatively limited number of samples (a total of 20 samples divided into two endpoint categories). This approach could be feasibly integrated into a biomarker strategy in early phase trials, deriving predictive biomarkers from a small number of patients and subsequently applying them in later phase trials to maximize likelihood of success.

## DATA AVAILABILITY

The single cell transcriptomics data will be released publicly upon manuscript publication.

## DISCLOSURE STATEMENT

Michael Tsabar, Sarah Sturtevant, Ryan Genga, Margaret Magaletta, Matthew Kodrasov, Kyle Smith, Andre Kurlovs, Bruno Figueroa, Alexandra Grella, Emanuele de Rinaldis, and Giorgio Gaglia are employed by Sanofi. Yinyin Huang, Elizabeth Tran, Juhyung Jung, Matthew Sullivan, and Virginia Savova are former Sanofi employee and may hold Sanofi stock. Shaoline Sheppard, Diana Stoycheva, and Dennis Goehlsdorf are Scailyte AG employees. Virginia Savova is a Scientific Advisory Board member of Scailyte AG.

### ACKNOWLEDGMENTS

Human blood sample used in the research were obtained from Oklahoma Blood Institute and Charles River Laboratories, then banked and processed by Sanofi CMC laboratories. The research was funded by Sanofi.

## MATERIAL AND METHODS

### Experimental and Laboratory methods

#### PBMC Sourcing

Peripheral leukapheresis donations from healthy individuals were obtained from Charles River Laboratories or Oklahoma Blood Institute. All donors were screened for CMV positivity and infectious disease panel. Collections are shipped at 4°C overnight for next day processing or were cryopreserved at the collection site and then shipped and stored in liquid nitrogen (LN2) for later use.

#### PBMC Processing, In-Process Analysis, with CD3 and/or CD19 Depletion

The CliniMACS Prodigy Automated Cell Processing System is utilized for GMP-compliant expansion of NK cells. On day 0, cryopreserved leukapheresis packs undergo a controlled thaw using a Plasmatherm. For every 3E9 nucleated cells available, thawed material is diluted with 70 mL of 37°C CliniMACS PBS/EDTA supplemented with HSA to 1%. Diluted starting material is passed through a Pall Blood Transfusion Filter and loaded into the Prodigy Centricult Unit (CCU) of a TS520 tubing set. PBMCs are then further washed with PBS/EDTA buffer supplemented with HSA to 0.5% and centrifuged to remove platelet content. If fresh leukapheresis material is being processed, density gradient separation (DGS) is performed with Ficoll-Paque to remove red blood cells.

With fresh leukapheresis, a sample is acquired from the CCU post-DGS. Total cell counts are determined by flow cytometry. Cells in the CCU are blocked by addition of Intravenous Immunoglobulin (IVIG) to 0.125 %. Cells are then incubated with anti-CD3, with or without anti-CD19, iron-conjugated microbeads for 30 minutes. Post-incubation, cellular material is sequentially passed over the magnetic column in the TS520 tubing set, 600E6 CD3+ cells at a time. The total number of column passes is determined by the in-process analysis described below. During the depletion, PBMCs are transferred to an external cell bag on the TS520 tubing set. The column and CCU are washed with sterile water and then PBS/EDTA/HSA buffer to eliminate any remaining T-Cells. The depleted PBMCs are passed over the column again to capture residual CD3+ cells and to transfer the cells back into the CCU. At the end of day 0 processing, NK cells were frozen down in CryoStor10 (CS10) at 5e6 NK cells/mL. One vial was labeled for sequencing and the other vial was saved for expansion.

#### T-flask Expansion of NK Cells

CD3 Depleted PBMCs that were cryopreserved post day 0 processing on the Prodigy in CryoStor10 were thawed using the ViaStar for PM21 stimulated expansion. The vials of CD3 depleted PBMCs was thawed, counted by flow cytometry, and stimulated with PM21 at 0.25 mg/mL, before being diluted to 0.3E6 NK cells/mL in cSCGM supplemented with 100 IU/mL of IL-2. On Day 4 of the process, an 80% media exchange is performed with cSCGM supplemented with 100 IU/mL IL-2.

On process days 5 and 6, cells were counted by flow cytometry and underwent 80% media exchanges. On the morning of day 7, a sample is pulled for counting via flow cytometry and the cells were restimulated with PM21 for 6 hours, before being diluted to 0.42E6 NK cells/mL in cRPMI, as previously described. For days 8-13, cells were counted by flow cytometry and diluted to 0.42E6 until a maximum volume of 50 mL was reached. At maximum volume, 100% media exchanges were performed. On Day 13, cells were counted by flow cytometry, and cytotoxicity, surface marker, and characterization panels were completed.

#### In-process monitoring of NK Cell Count and Viability

Two 100 μL samples of wild type process samples were obtained and incubated with 5 μL of TruStain FcX Receptor Blocking solution for 5 minutes. Next, one set of cells were stained with 120 ng CD56 PE and 75 ng CD3 APC for 8-10 minutes, while the isotype control was treated with 120 ng IgG1, κ PE and 75 ng IgG1, κ APC and incubated for the same duration. After incubation, each sample was treated with 0.3 μM DRAQ7 for 2 - 3 minutes. Stained samples were transferred to TruCount Absolute Counting Tubes. Samples were then acquired on the FACS Lyric.

#### CITE-seq data generation

Starter cultures from donor blood were generated as described above and samples were cryopreserved and stored at –80 °C. Frozencells were thawed dropwise into pre-warmed RPMI-1640 medium supplemented with 10% fetal bovine serum. Cells were then washed twice and counted using the Cellaca MX cell counter (Revvity, USA). For surface protein labeling, cells were stained using the TotalSeq™-B Human Universal Cocktail (BioLegend, Cat# 399904). Approximately 500,000 cells were resuspended in cold phosphate-buffered saline (PBS) containing 0.05% bovine serum albumin (BSA). Fc receptors were blocked by incubating cells for 10 minutes with TruStain FcX™ (BioLegend, USA) to minimize nonspecific antibody binding. Cells were subsequently incubated with barcoded antibody mixtures for 30 minutes at 4 °C. Following incubation, cells were washed three times with PBS + 0.05% BSA via centrifugation (~300 × g for 5 minutes at 4 °C). After the final wash, cells were resuspended at the appropriate concentration in PBS for downstream 10x Genomics applications.

#### Single-Cell RNA Sequencing

Single-cell capture and library preparation were performed using the Chromium Next GEM Single Cell 3’ Reagent Kits v3.1 (Dual Index) and Feature Barcode technology for Cell Surface Protein Labeling (10x Genomics, USA), following the manufacturer’s protocols (CG000149 and CG000317 Rev D). Final libraries were pooled and sequenced on the Illumina NovaSeq 6000 platform.

### Data processing and analysis

#### CITE-seq preprocessing and statistical analysis pipeline

From raw sequencing data, UMI count matrices were generated by Cellranger V5.0.0. The single-cell processing pipeline CellBridge [27], was used to perform pre-processing from HDF5 format to a packaged Seurat object. CellBridge, in brief, translates and packages HDF5 files into R-based Seurat object and performs QC thresholding (minimum UMI = 750, percent.mt < 20, nFeature_RNA<250), a log normalization of count data (LogNormalize function with a scaling factor of 1e+06), FindVariableFeatures (nfeatures=2000), RunUMAP (dims = 1:30, res=0.7, k=20), and Harmony (by sample). Automated celltype calling was performed by SignacX [28], a ML-based classification algorithm.

Primary data visualization pipelines in R were run under the R 4.3.2 environment. UMAPs were plotted using native Seurat (5.1.0) functions. Cell type proportion comparisons were visualized using the ggboxplot function in the ggpubr (0.6.0) package and significance comparisons were performed using an unpaired t-test for the cell types considered in the plot. For differential expression analyses between conditions, a custom wrapper function using edgeR (4.0.16)/limma (3.58.1) was created. In brief, this function performs pseudobulk aggregation using the function AggregateExpression on a Seurat object’s raw count data, per condition_cellstate. Then, we created a pairwise design and contrast matrix and the equation ~0+comparsioncellstates with no input covariates into the analyses. A standard recommended pipeline for creating a negative binomial generalized linear model with F-test using filterByExpr, calNormFactors, estimateDisp, and glmQLFit was then called and individual glmQLFTests were run in parallel using foreach (1.5.2) and doParallel (1.0.17) packages. Differential expression results were visualized using the EnhancedVolcano (1.14.0) package.

#### ScaiVision modeling

Prediction of NK cell expansion was conducted using the neural network algorithm CellCnn (Eirini Arvaniti, Manfred Classen, 2017), implemented in PyTorch in the ScaiVision platform (Scailyte AG, version 1.6.4), similar to Roussel et al. ScaiVision is a supervised machine learning algorithm that trains a convolutional neural network with a single hidden layer on single-cell data to predict sample-level outcomes. Top and lowest 10 samples based on ex vivo NK cell counts recorded after expansion were extracted and relabeled as high expansion and low expansion, respectively. Samples with intermediate cell counts were excluded from model training. A 3-fold Monte Carlo cross-validation scheme with a 30/70 split was employed to evaluate the reproducibility of the performance. Monte-Carlo sampling was chosen over a leave-on-out-scheme because the validation set is also used to monitor training and best model selection. For this process, several samples are required to compute a meaningful selection criterion. The gene-by-cell matrix was first reduced to principal components to minimize overfitting by lowering the number of trainable parameters. For model training using all cells, 30 components were used for gene expression models and 20 for models using surface protein abundance. After selecting specific cell types, the number of components was adjusted: 10 components were used for NK and B cells individually, and 20 components were used when both cell types were selected. The dimension reduction step was excluded for training runs with restricted gene sets of 100 or fewer genes. For each sample, sub-samples of 500 cells, called multi-cell inputs (MCIs), were randomly selected. For each training epoch, 1000 MCIs per class (High NK cell expansion vs Low NK cell expansion) were presented to the network in random order. A total of 50 independent networks were trained using hyperparameters randomly chosen from the following options: i) number of filters: 3, 5, and 10, ii) top-k pooling percentage: 2, 5, 10, 20, iii) dropout probability: 0.3, 0.4, and 0.6, iv) learning rate: 0.0003, 0.0001, 0.001, 0.003 and 0.01, and v) weight decay: 0.00001, 0.0001, 0.001, 0.01, and 0.1. Training was performed with a batch size of 64. Each network was trained for a maximum of 50 epochs, or until the validation loss no longer decreased for 20 consecutive epochs. Models were eventually evaluated using the parameters restored from the training epoch which yielded the lowest validation loss.

#### Extraction of predictive cells

The results of the ScaiVision modeling consist of both a trained model capable of predicting the outcome of interest as well as the molecular characterization of the most relevant cells associated with that outcome. For each CV-split, models with a validation accuracy above 0.8 were selected. By multiplying the weights of each filter with the gene expression matrices through a weighted sum, a score per filter and cell is obtained. This score represents how strongly the cells respond to the filter. To identify predictive cells across filters, cells were ranked for each filter and the average rank across filters was calculated. The top 3% of cells with the highest score were annotated as predictive cells.

#### Calculation of gene contribution scores

Models were analyzed using the Captum library (Sundararajan, Taly, Yan, 2017). For each CV-split, models with a training and validation accuracy of ≥0.75 were shortlisted, and the top 3 based on validation loss were selected. Validation loss was picked as a sorting criterium because it allows to better differentiate good models and favors models yielding higher separation between samples of different classes. Feature importance scores were computed using deepLIFT (Shrikumar, Greenside, Kundaje, 2019) and integrated gradients (Proceedings of the 34th International Conference on Machine Learning, 2017). These methods each provide a metric for how each feature contributes to predicting high or low NK cell expansion. Scores were calculated on models trimmed off from their final SoftMax layer to produce log-space values, enabling convenient addition for combination. Each algorithm generates contribution values for each combination of model, gene, cell, and prediction neuron. These values were averaged across cells within each sample, and the difference between the two output neurons’ averages was calculated. This yielded a score reflecting how strongly a specific gene drove the prediction of high NK cell expansion for each sample. Scores were grouped by actual high or low expansion samples, averaged within each group, and the difference was used to assess gene contribution accuracy. Finally, these contribution scores were averaged across the selected models.

## SUPPLEMENTARY INFORMATION

**Figure S1:**
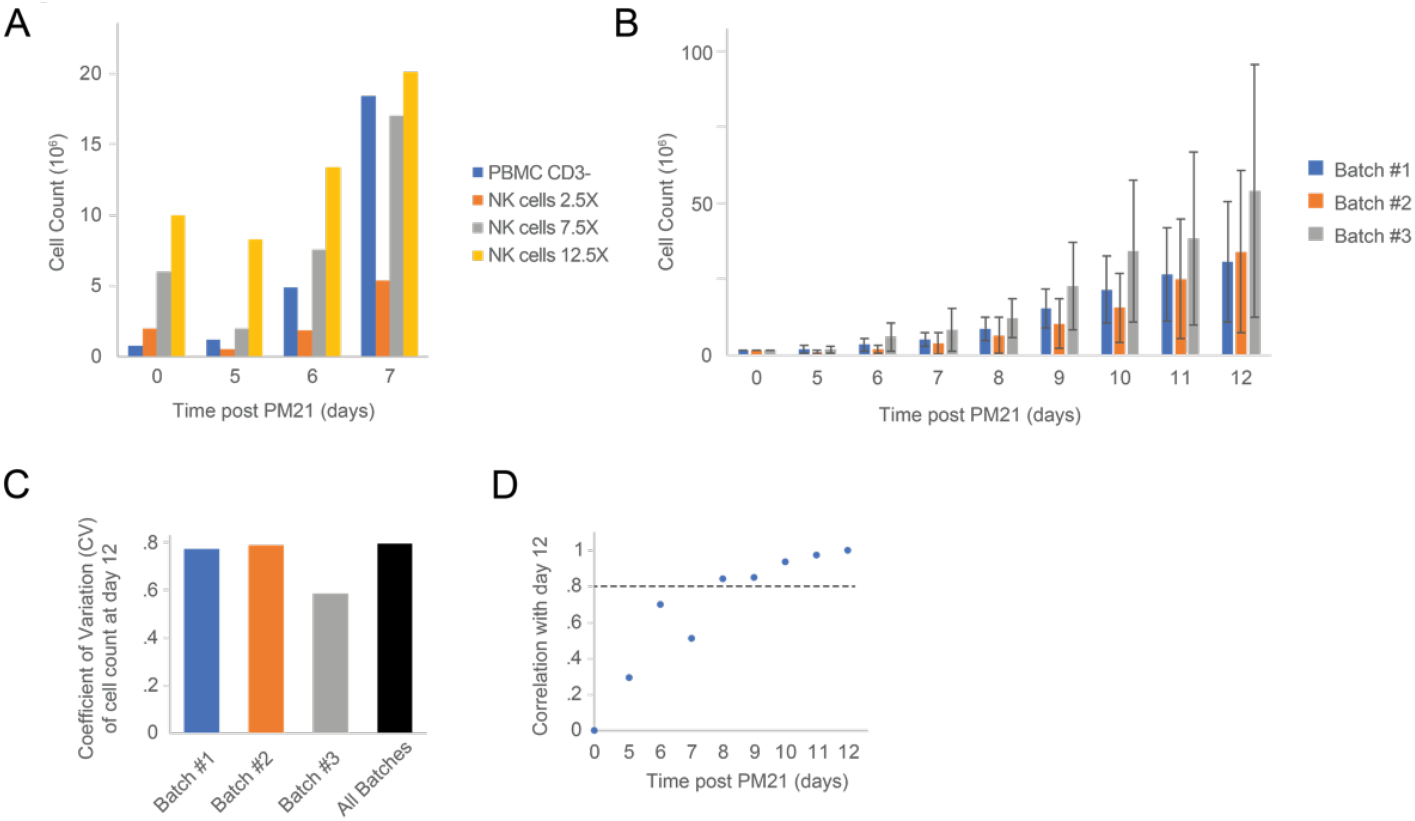
**A**. Comparison of expansion rate for T-cell depleted PBMC’s (PBMC CD3-, blue), and isolated NK cells at increasing relative numbers compare to PBMC CD3-(2.5-fold, orange, 7.5-fold, grey, and 12.5-fold, yellow) from same donor (n=1). **B**. Expansion rate by batch (mean +/− standard deviation, n=9,8,9 per batch). **C**. Coefficient of variation across expansion batches #1-3 (n=9,8,9), and for all samples together (“All Batches”, n=26, black bar). **D**. Pearson correlation of NK cell counts at endpoint (day 12) and earlier timepoints (n=26 donors). Dotted grey line y=0.8 for comparison.

**Figure S2:**
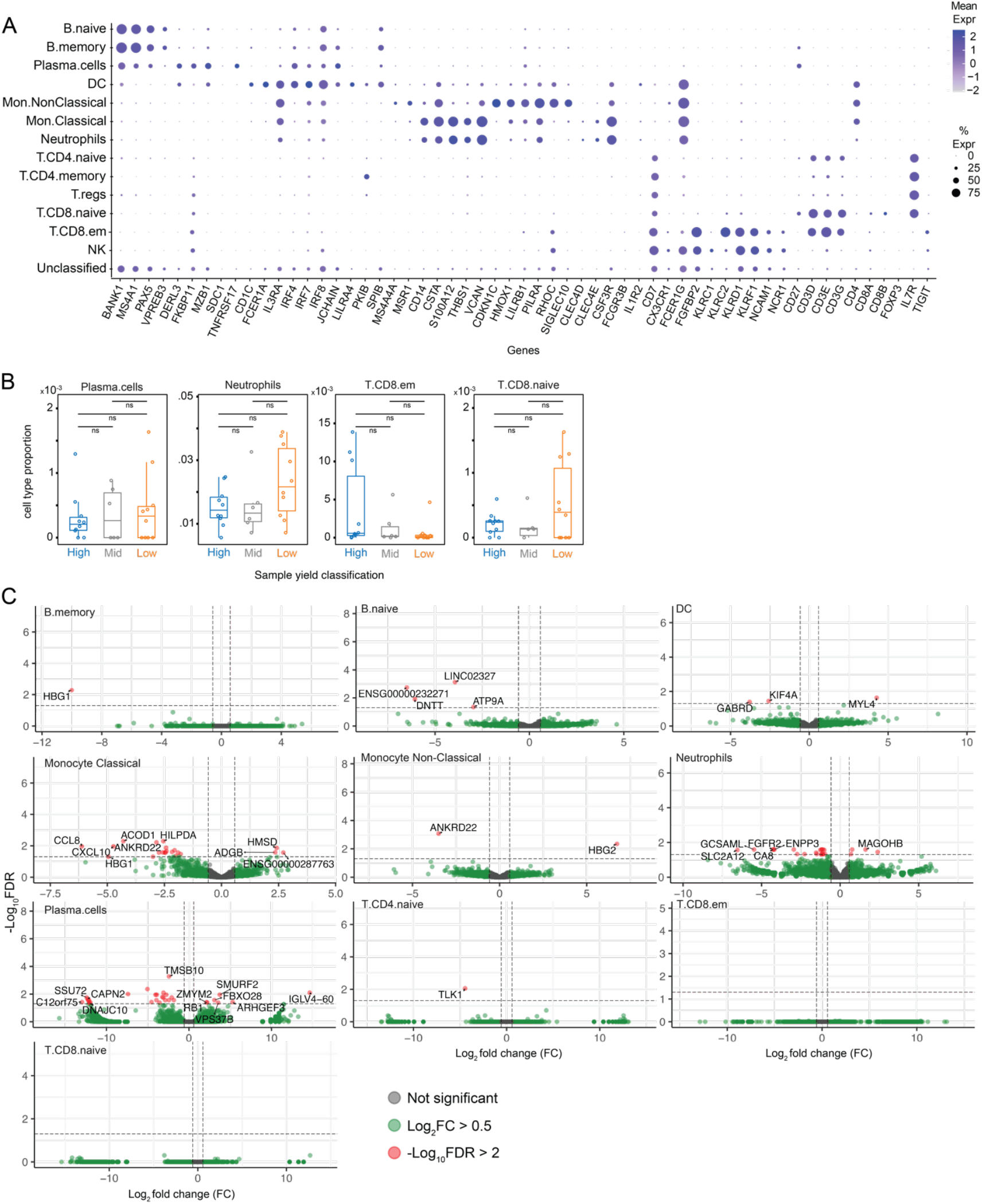
**A**. Seurat bubble plot of canonical cell type markers by cell types defined by SignacX. **B**. Box plot with whiskers of SignacX cell state proportions subdivided by high-, medium- and low-yield expanders (High, n=10, blue, Mid n=6, grey, Low n=10, orange). **C**. Volcano plot of pseudobulk differential gene expression of per cell type in high-versus low-yield expanders.

**Figure S3:**
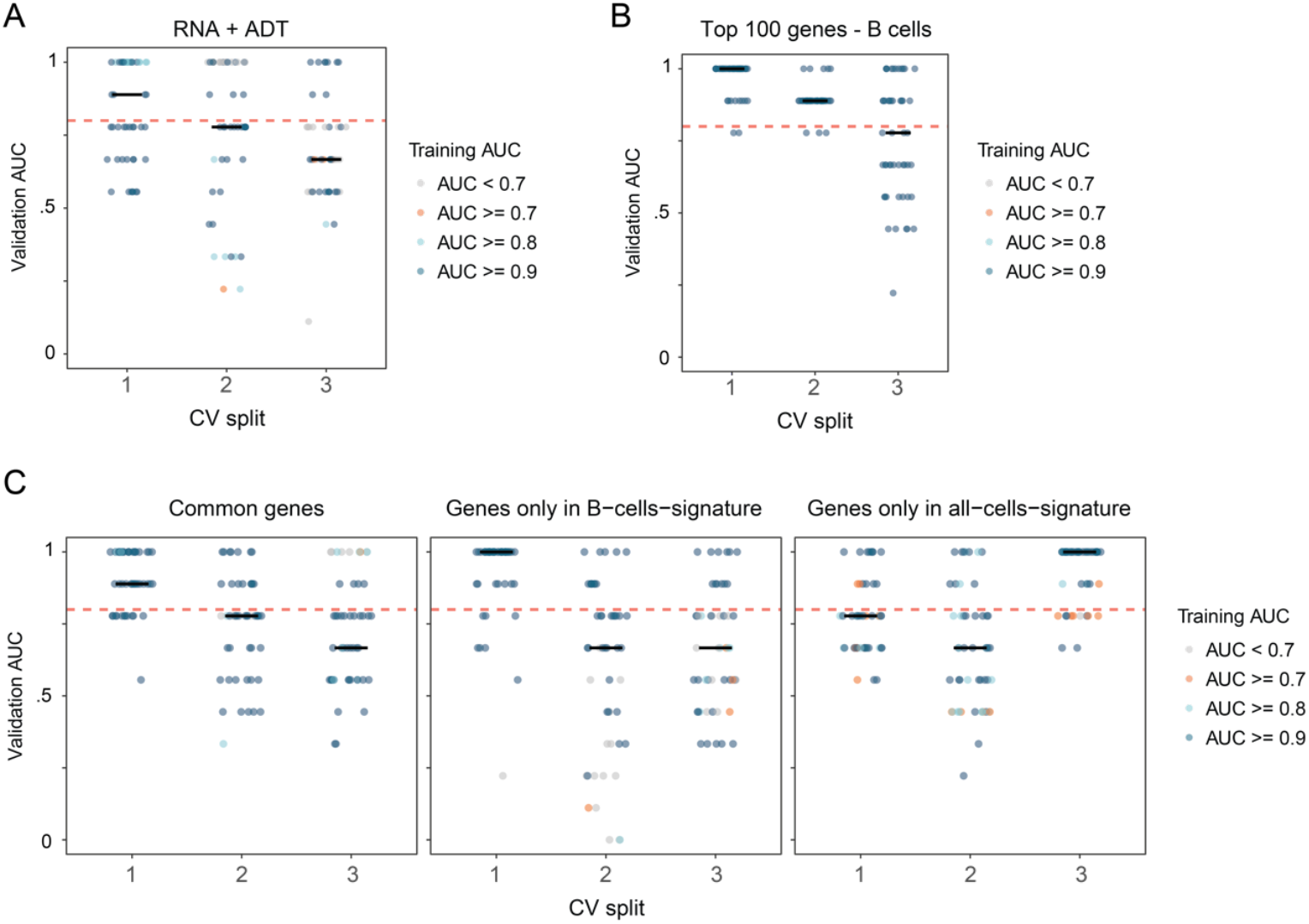
**A**. Area under the curve (AUC) from validation of n=50 models per CV-split for models utilizing the combined ADT and RNA data. **B**. Area under the curve (AUC) from validation of n=50 models per CV-split. Models were trained only using B cells using the top 100 genes derived from B-cell models. **C**. Area under the curve (AUC) from validation of n=50 models per CV-split. Models were trained only using either the common genes between the B-cell and all-cell top 100 genes (left), the genes exclusive to the B-cell top 100 gene set (center), or the genes exclusive to the all-cell top 100 gene set (right). In all panels, color represents AUC achieved by each model in training phase. The horizontal black line represents median AUC.

**Supplementary Table S1:** Sample cross-validation split.

| Sample ID | NK cell counts | Expansion categories | Batch | CVsplit1 | CVsplit2 | CVsplit3 |
| --- | --- | --- | --- | --- | --- | --- |
| S49 | 1290000 | low | 1 | Test | Test | Training |
| S37 | 5210000 | low | 2 | Training | Training | Training |
| S28 | 8580000 | low | 3 | Test | Training | Test |
| S40 | 9090000 | low | 2 | Training | Training | Test |
| S27 | 10900000 | low | 3 | Test | Training | Training |
| S31 | 13900000 | low | 3 | Training | Training | Training |
| S41 | 18000000 | low | 2 | Training | Test | Training |
| S50 | 19300000 | low | 1 | Training | Test | Test |
| S44 | 20000000 | low | 1 | Training | Training | Training |
| S38 | 22600000 | low | 2 | Training | Training | Training |
| S35 | 24500000 | medium | 3 | Validation | Validation | Validation |
| S42 | 25400000 | medium | 2 | Validation | Validation | Validation |
| S33 | 25700000 | medium | 3 | Validation | Validation | Validation |
| S29 | 32400000 | medium | 3 | Validation | Validation | Validation |
| S32 | 37400000 | medium | 3 | Validation | Validation | Validation |
| S48 | 38600000 | medium | 1 | Validation | Validation | Validation |
| S52 | 43700000 | high | 1 | Training | Test | Training |
| S45 | 51700000 | high | 1 | Training | Test | Test |
| S36 | 53200000 | high | 2 | Test | Training | Training |
| S30 | 55500000 | high | 3 | Training | Training | Training |
| S43 | 57000000 | high | 2 | Training | Training | Training |
| S34 | 66200000 | high | 3 | Training | Training | Training |
| S47 | 77800000 | high | 1 | Test | Test | Training |
| S39 | 80500000 | high | 2 | Training | Training | Test |
| S51 | 111000000 | high | 1 | Test | Training | Test |
| S46 | 121000000 | high | 1 | Training | Training | Training |

**Supplementary Table S2:** Summary of predictive cell enrichment.

| cell_type | cell_type.size | fraction_predictive_cells | hg.pval | hg.FDR | cvsplit |
| --- | --- | --- | --- | --- | --- |
| B.memory | 7989 | 0.19739642 | 0 | 0 | cvsplit1 |
| B.memory | 7989 | 0.159093754 | 0 | 0 | cvsplit2 |
| B cells | 27971 | 0.12162597 | 0 | 0 | cvsplit1 |
| B cells | 27971 | 0.104000572 | 0 | 0 | cvsplit2 |
| B.naive | 19955 | 0.091054873 | 9.67E-298 | 5.80E-297 | cvsplit1 |
| B.naive | 19955 | 0.081934352 | 2.10E-208 | 1.26E-207 | cvsplit2 |
| T cells | 452 | 0.504424779 | 2.24E-194 | 1.01E-193 | cvsplit2 |
| T cells | 452 | 0.504424779 | 2.24E-194 | 1.05E-193 | cvsplit2 |
| B.memory | 7989 | 0.105770434 | 1.57E-158 | 2.83E-157 | cvsplit3 |
| DC | 4138 | 0.121072982 | 1.78E-113 | 6.22E-113 | cvsplit2 |
| DC | 4138 | 0.121072982 | 1.78E-113 | 6.40E-113 | cvsplit2 |
| T cells | 452 | 0.314159292 | 4.98E-87 | 2.24E-86 | cvsplit1 |
| T cells | 452 | 0.314159292 | 4.98E-87 | 2.33E-86 | cvsplit1 |
| B cells | 27971 | 0.059168424 | 1.63E-77 | 2.29E-76 | cvsplit3 |
| NK | 18555 | 0.063702506 | 1.94E-69 | 6.81E-69 | cvsplit1 |
| NK | 18555 | 0.063702506 | 1.94E-69 | 7.00E-69 | cvsplit1 |
| NK | 18555 | 0.052061439 | 1.06E-21 | 2.97E-21 | cvsplit2 |
| NK | 18555 | 0.052061439 | 1.06E-21 | 3.18E-21 | cvsplit2 |
| Neutrophils | 1506 | 0.090305445 | 2.42E-19 | 8.47E-19 | cvsplit3 |
| Neutrophils | 1506 | 0.090305445 | 2.42E-19 | 1.09E-18 | cvsplit3 |
| Plasma.cells | 27 | 0.296296296 | 5.19E-07 | 1.17E-06 | cvsplit1 |
| Plasma.cells | 27 | 0.222222222 | 6.34E-05 | 0.000190326 | cvsplit3 |
| Plasma.cells | 27 | 0.111111111 | 0.020007817 | 0.045017587 | cvsplit2 |
| B.naive | 19955 | 0.040290654 | 0.199406452 | 0.448664517 | cvsplit3 |
| DC | 4138 | 0.01377477 | 1 | 1 | cvsplit1 |
| Monocytes | 71525 | 0.000894792 | 1 | 1 | cvsplit1 |
| Neutrophils | 1506 | 0.003320053 | 1 | 1 | cvsplit1 |
| Monocytes | 71525 | 0.003495281 | 1 | 1 | cvsplit2 |
| Neutrophils | 1506 | 0.005976096 | 1 | 1 | cvsplit2 |
| DC | 4138 | 0.010874819 | 1 | 1 | cvsplit3 |
| Monocytes | 71525 | 0.037273681 | 0.999983705 | 1 | cvsplit3 |
| NK | 18555 | 0.01956346 | 1 | 1 | cvsplit3 |
| T cells | 452 | 0.019911504 | 0.984074771 | 1 | cvsplit3 |
| DC | 4138 | 0.01377477 | 1 | 1 | cvsplit1 |
| Monocytes | 71525 | 0.000894792 | 1 | 1 | cvsplit1 |
| Neutrophils | 1506 | 0.003320053 | 1 | 1 | cvsplit1 |
| Monocytes | 71525 | 0.003495281 | 1 | 1 | cvsplit2 |
| Neutrophils | 1506 | 0.005976096 | 1 | 1 | cvsplit2 |
| DC | 4138 | 0.010874819 | 1 | 1 | cvsplit3 |
| Monocytes | 71525 | 0.037273681 | 0.999983705 | 1 | cvsplit3 |
| NK | 18555 | 0.01956346 | 1 | 1 | cvsplit3 |
| T cells | 452 | 0.019911504 | 0.984074771 | 1 | cvsplit3 |

